# Shifts and rebound in microbial community function following repeated introduction of a novel species

**DOI:** 10.1101/2024.04.05.588252

**Authors:** Alanna M. Leale, Francisca Reyes Marquez, Bas Zwaan, Eddy J. Smid, Sijmen Schoustra

## Abstract

Natural microbial communities continually encounter novel species that may successfully establish or simply be transient, yet both outcomes can alter the resident community composition and function. Preserving natural microbial communities and innovating synthetic ones requires insight on the immediate and long-term impact of species introductions on both composition and function. For instance, it remains unclear whether there are gradual and long-term impacts from repeated invasions where the introduced species fails to establish – so-called failed invaders. To investigate the persistent impacts by failed invaders, we present an experimental test of community stability over multiple generations against repeated novel species introduction. We propagated a natural microbial community from a traditional fermented milk beverage for approximately 100 generations, with or without, repeated introduction of *Escherichia coli* at each transfer. Community function was determined by metabolic profiling, and we observed alterations therein immediately after *E. coli* introduction, followed by recovery, or rebound once ceased. In contrast to this proxy of community function, changes in the bacterial community composition were never detected. Our results evidence that community composition and function do not necessarily respond in parallel to an introduced species, potentially due to genotypic changes below species level detection or metabolic plasticity. Our work shows an ability for functional recovery in microbial communities and contributes insight on long-term community stability to sustained disturbances.

## INTRODUCTION

Natural ecosystems and the species communities they harbour are commonly faced with perturbations by invading novel species, whether gradual and continuous or in singular events. Thinking of invasive species typically recalls non-native plants, insects, or animals proliferating in a habitat. For instance, zebra mussels in North American fresh lakes (Karatayev and Burlakova 2022), the common water hyacinth plant in tropical ponds globally (Datta *et al*. 2021), or the emerald ash borer beetle in European and North American forests (McCullough 2020), are just a few well-known cases when introduced species successfully, or devastatingly *per se*, established in a novel environment. The same processes occurring at a macroecological level also apply to microbial ecosystems that harbour communities of co-existing microbes. The spoilage of food (Oro *et al*. 2019), cyanobacterial blooms (Bolius, Wiedner and Weithoff 2019), biofertilizers (Gu *et al*. 2019), probiotics (Albright *et al*. 2020), or infection of human microbiota (Libertucci and Young 2019), can all be considered instances where the resident microbial community is confronted by novel species (Kinnunen *et al*. 2016). But even if unsuccessful at establishing, an introduced species can change the structure of interactions between members of a microbial community (Padrón *et al*. 2009) and furthermore influence overall community function (Mallon *et al*. 2018), even shifting the community to an alternative stable compositional state (Amor, Ratzke and Gore 2020). Further, community species composition is found to be decoupled from community metabolic function (Louca *et al*. 2016, 2018) meaning that responses to perturbation at one level may not align with the other. As the great value of microbial communities for both natural ecosystems and human-desired functions is rapidly being recognised (Kinnunen *et al*. 2016; Sivasubramaniam and Franks 2016), it is of interest to know how they may respond to perturbations and how responses in community composition *versus* function contrast, including the persistence of shifts to alternative states once perturbation stops.

Protecting natural microbial communities or engineering novel synthetic ones both require understanding immediate and long-term responses of community composition and function to perturbation by novel introduced species. The stability of microbial communities against perturbation by a novel species has been investigated with multiple experimental and modelling approaches across study systems (Mallon, Elsas and Salles 2015; Vila *et al*. 2019; Kruk *et al*. 2021; Kurkjian, Akbari and Momeni 2021; Philippot, Griffiths and Langenheder 2021). Much research has focused on a single introduction of a novel species to microbial communities over a few generations of a single growth cycle (Allison and Martiny 2008; Shade *et al*. 2012; van Elsas *et al*. 2012; Mallon *et al*. 2018; Amor, Ratzke and Gore 2020), with few studies assessing multiple rounds of propagation and assessments at multiple time points (Xing *et al*. 2021; Albright *et al*. 2022). Single “flash” introduction events are undoubtably important to understand as they occur in natural ecosystems; however, it is also common that novel species are repeatedly introduced into a community with compound effects (Blackburn, Lockwood and Cassey 2015; Philippot, Griffiths and Langenheder 2021). Specifically, a key finding motivating our study is that even a transient novel species, or a failed invader, can induce a lasting alteration to a bacterial community’s composition and resource use breadth (Yao *et al*. 2014; Mallon *et al*. 2018; Amor, Ratzke and Gore 2020; Xing *et al*. 2021). The permanence of induced changes is relevant to investigate – do microbial communities rebound in metabolic function and composition, or remain altered after repeated exposure to a failed invader? To test this, observations on community stability against novel species over extended timescales of multiple generations are needed (Philippot, Griffiths and Langenheder 2021; Albright *et al*. 2022), and would help elucidate unanswered questions about the persistent impacts by failed invaders.

We used a microbial community of a traditionally fermented milk beverage from Zambia – Mabisi – as a model system to investigate the impact of repeated introduction of a novel species on community function. Microbial communities of Mabisi sampled in the field typically contain a diverse yet manageable composition of approximately 6 to 12 species of bacteria, but samples can vary depending on location and processing methods (Schoustra *et al*. 2013; Moonga *et al*. 2020). As an introduced species, we chose a non-pathogenic *E. coli* strain for its relevance to food safety in spontaneously fermented foods (Capozzi *et al*. 2017) and ease of laboratory use for selective plating at a low biohazard level. *E. coli* is also used in other microbial community stability research (van Elsas *et al*. 2012; Mallon *et al*. 2018; Xing *et al*. 2021) due to its ability to grow, or at least persist, in secondary environments from its primary environment of the vertebrate gut (Savageau 1983; van Elsas *et al*. 2007). Mabisi is traditionally made by spontaneous fermentation of raw, unpasteurised milk, thus its microbiological safety is called into question and its sale restricted to household level in Zambia (Materia *et al*. 2021). However, past research exhibited that Mabisi communities are highly resistant against food borne pathogens including *Enterobacteriaceae* (Schoustra *et al*. 2022); we therefore did not predict *E. coli* establishment. Expected failed *E. coli* invasion in Mabisi therefore provided a suitable approach to evaluate how the resident community species composition and function would be impacted over repeated failed invasion events, as well as community recovery afterwards.

The varying macro-ecological and microbial systems for researching ecological stability has led to inconsistent terminology (Donohue *et al*. 2016; Kinnunen *et al*. 2016; Philippot, Griffiths and Langenheder 2021), so we here stablish our definitions. We refer to the resident community as the members and their relative abundances stably present in a natural setting, whereas the novel species as one that is absent or at negligible abundances in the initial resident community, hence considered introduced. We do not use the term “invader” due to the term’s implication of domination, proliferation, or at least establishment, whereas simply the alteration of community structure, regardless of relative abundance (i.e., establishment) of the novel species, is of focus in our work. In line with other microbial ecology studies on stability, our study assessed both resistance and recovery in a microbial community (Allison and Martiny 2008; Shade *et al*. 2012; Philippot, Griffiths and Langenheder 2021), doing so in terms of metabolic function and community composition (Table 1). Resistance is the degree of change in a variable, for example, how much does the community composition or function shift following disturbance. Recovery is if, and the degree that, the community returns to its pre-disturbance state (Shade *et al*. 2012). Metabolic profiles are an especially suitable proxy for community function in fermented foods since they indicate compounds associated with aroma and taste (Alekseeva *et al*. 2021), while inferring to active pathways and gene expression across the community (Smid *et al*. 2005; De Filippis *et al*. 2016). Acidity was additionally measured since it influences consumer preference (Ott *et al*. 2000; Sikombe *et al*. 2023) but more importantly, the prevention of establishment of food borne pathogens in fermented foods (Mpofu *et al*. 2016).

**Table 1:** community stability related definitions and measures, as used in this paper.

| Term | Definition related to this study’s results |
| --- | --- |
| resident community | composition of community members stably present in a natural setting |
| introduced/novel species | species that is absent or at negligible abundances in the initial resident community |
| resistance | change in metabolic function and community composition following disturbance. Little or no change = high resistance. |
| recovery | return to a pre-disturbance state of metabolic function and community composition once disturbance is stopped. |
| community composition | full-length 16s rRNA gene sequences clustered at 95% similarity |
| community function | metabolic profiles (GC-MS analyses) and acidity |

In this study, our general research aim was to investigate repeated introduction of a novel species to a microbial community over many generations. Our experimental approach had two parts: we first tested if, when repeatedly introduced, *E. coli* is unable to establish, but still can shift the Mabisi resident community composition and function? Secondly, we were interested in the Mabisi resident community’s recovery to rebound in composition and function once introduction was stopped. To do so, we propagated replicate natural Mabisi communities for 18 growth cycles (approximately 110 mitotic generations) in the laboratory under two treatments; a “Control” treatment without the addition of non-pathogenic *E. coli*, and an “Introduction” treatment where *E. coli* was added at every transfer (Figure 1). After 14 transfers, the “Introduction” treatment replicates were then split to generate a third “Recovery” treatment where *E. coli* was no longer added for the remaining transfers. Community composition (species identification through full 16s rRNA gene) and metabolic profiles (GC-MS) were measured throughout to assess the community resistance and recovery to repeated novel species introduction.

**Figure 1:**
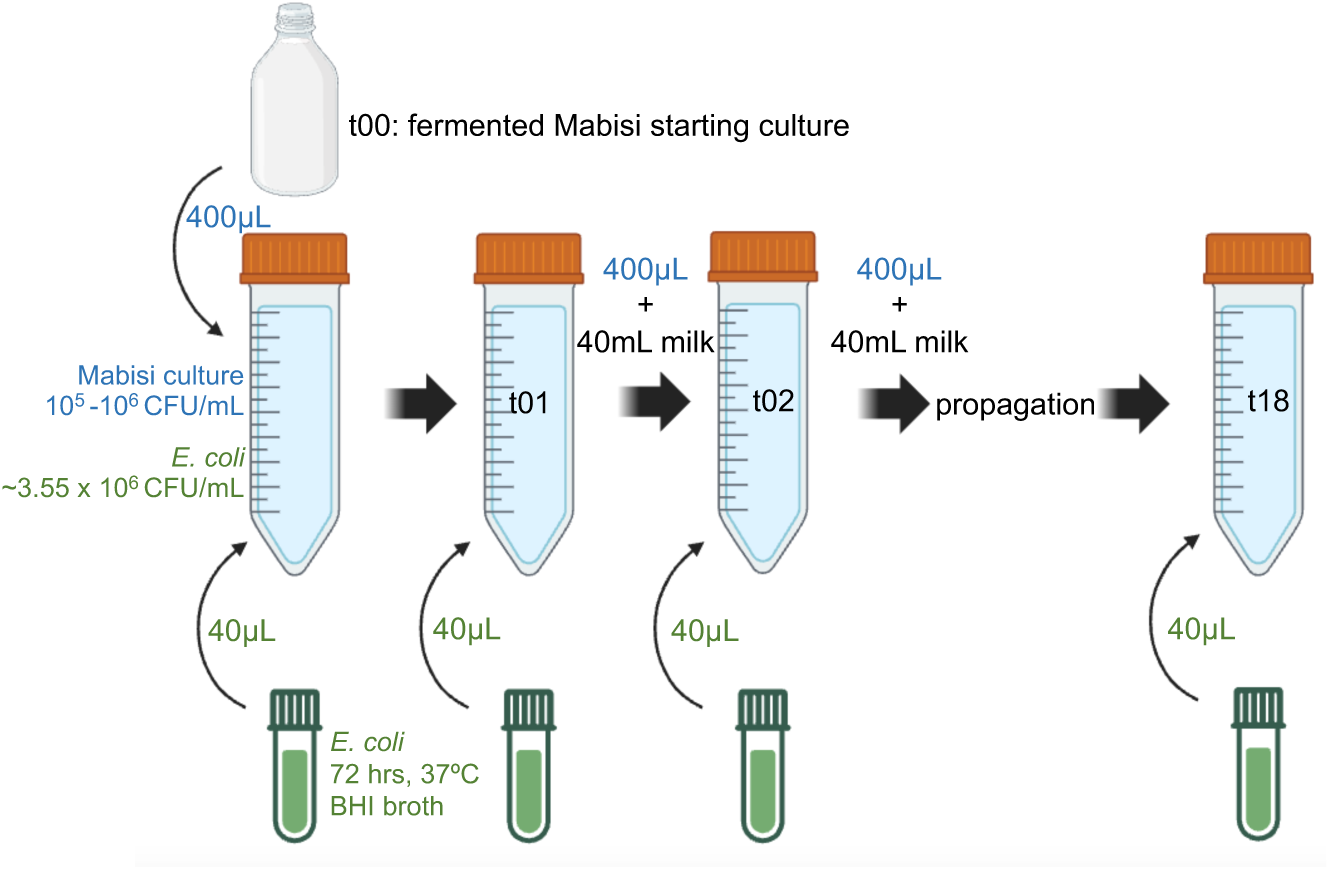
experimental set up of “Introduction” treatment. “Control” treatment was identical except without addition of *E. coli*. Both treatments were performed in replicates of six. At transfer 14, the “Introduction” replicates were used to create a “Recovery” treatment where *E. coli* was no longer added at each transfer. Please note that the same *E. coli* stock was used across transfers (i.e., *E. coli* was not evolving). Figure created in BioRender.

## RESULTS AND DISCUSSION

Using full-length 16s rRNA gene Oxford Nanopore sequencing at four time points, we assessed whether repeated *E. coli* introduction shifted resident bacterial community compositions and whether recovery was observed after introduction stopped. We were interested in the ecological process of species sorting – the sorting of variation at the level of species over time – in response to novel species introduction. We did not detect shifts in bacterial community composition between treatments of repeated *E. coli* introduction, at least at the full-length 16s rRNA sequencing level of bacterial type detection. The compositions of bacterial types did not differ between treatments but did change overtime (Table S1: PERMANOVA on community compositions, Bray method used: effect of treatment (p = 0.085, F = 2.78, DF = 2), transfer (p = 0.001, F = 16.83, DF = 3), diversity x transfer interaction (p = 0.034, F = 3.60, DF = 3)). Most notably, our starting Mabisi community contained lower bacterial type richness than expected (Table S2: mean species richness transfer 1 of Control treatment = 2.67, Invasion treatment = 3), based upon prior experiments using the same sample (Leale *et al*. 2023) we assumed to find approximately five types (three lactic acid bacterium types and two acetic acid bacterium types).

Although we did not detect changes in community composition at the resolution of the full-length 16s rRNA gene in the Introduction treatment (Fig. 2), it should not necessarily be inferred that underlying evolutionary processes are similarly static. So called “cryptic dynamics” has been proposed to describe the potential role of evolution in achieving ecological stability (Kinnison, Hairston Jr and Hendry 2015; Hendry 2019). In our experiment for example, there may be shifts in genotypes that are undetected since we can only detect types in the full-length 16s rRNA gene region, thus suggesting the community remains stable at the ecological level of bacterium type composition. Metagenomic sequencing would be required to investigate potential eco-evolutionary feedback mechanisms by detecting changes at the genotype, rather than species level. Although our null expectation of no change in community composition was observed, future research should look deeper into whether there are hidden evolutionary shifts in metabolism or gene expression.

**Figure 2:**
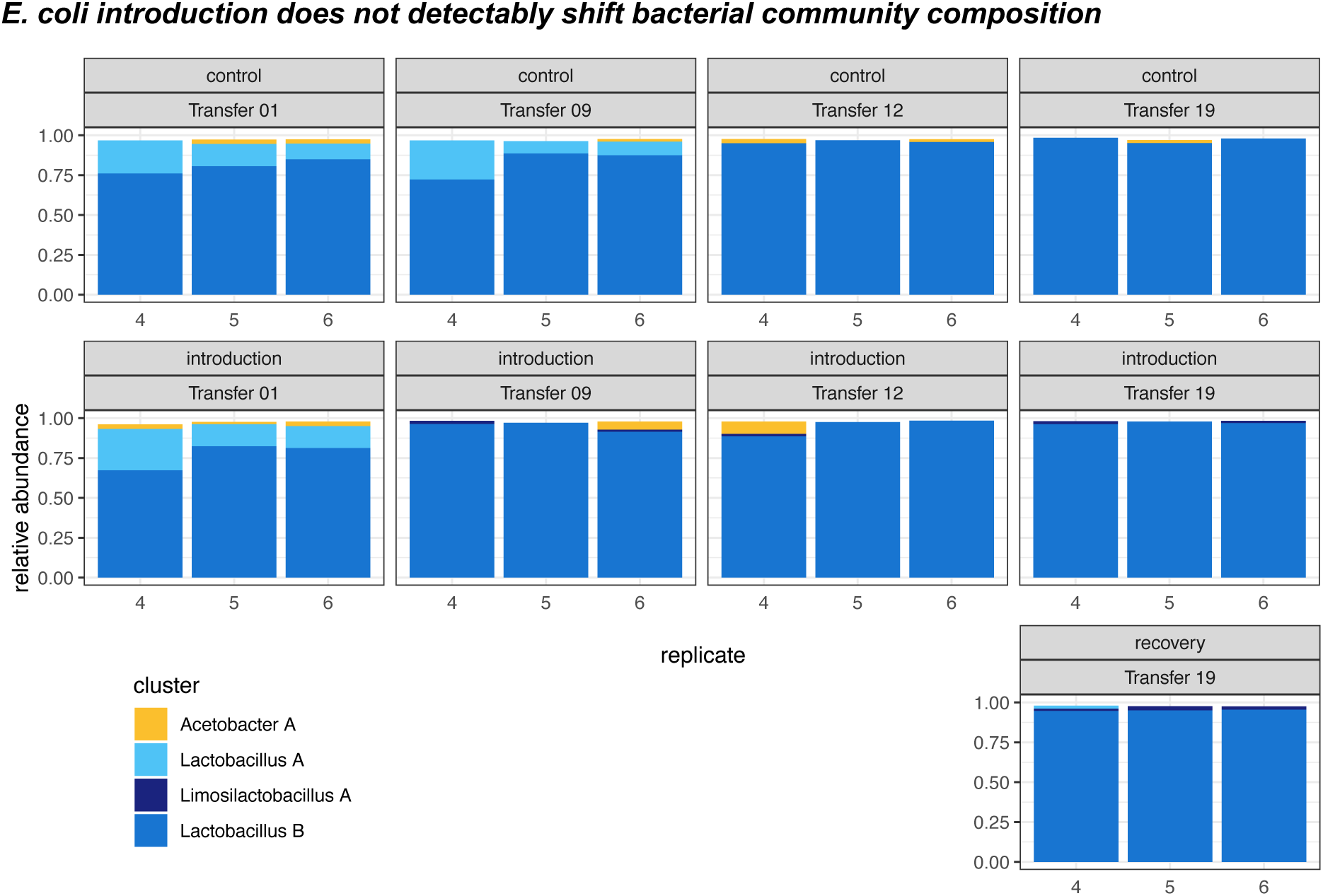
bacterial community profiles show no shift in species composition from E. coli introduction. Bacterial community compositions from full-length 16s rRNA sequencing at transfers 1, 9, 12, and 19. Only three of six replicates shown due to technical errors with sequencing in the first batch. Empty bars are clusters of <1% abundance which were not identified nor plotted. Transfer number refers to the end products of previous propagation (i.e., “transfer 9” are community compositions of the end products from transfer 8).

The full resistance of our Mabisi communities to *E. coli* establishment, at least to detectable levels after 72 hours of fermentation, was expected considering *E. coli’s* optimal pH of 6.5 to 7.5 (Davey 1994). Our results differ from others’ who observed *E. coli* successfully establish in new environments over a long period, such as soil water sheds over an entire year (Ishii *et al*. 2006). Although *E. coli* is known for impressive persistence and adaptation to new environments, where it has survived for up to seven days in soil pH of 4.57-5.14 (van Elsas *et al*. 2007; Zhang *et al*. 2013), this is still however above Mabisi’s final pH of approximately 4.0 or lower (Groenenboom *et al*. 2020; Moonga *et al*. 2020; Leale *et al*. 2023). The inability of *E. coli* to establish in our experiment even after 18 rounds of repeated introduction, strengthens the argument for Mabisi’s biological safety and upscaling beyond household production in Zambia (Schoustra *et al*. 2022).

Overall, we observe high resistance of Mabisi communities against *E. coli* in terms of resident bacterial community species composition and establishment. Unfortunately, the lack of species diversity in starting communities may have restricted our ability to detect potential changes in the composition of resident species. Furthermore, in the present study we focussed on potential shifts in the bacterial community and this approach overlooks potential changes in yeast abundances which are also present in the populations used here and our previous study (Leale *et al*. 2023). Regardless, we can say that the introduction of *E. coli* did not exert great enough pressure to induce bacterial species sorting, yet this was not true when considering community function responses, which we elaborate in the next section.

Resistance and recovery in community function in response to repeated novel species introduction was tested using the production of volatile organic compounds (VOCs) as a proxy (Smid *et al*. 2005; De Filippis *et al*. 2016; Alekseeva *et al*. 2021). By measuring metabolic profiles of VOCs overtime, we observe changes in community function in response to *E. coli* introduction immediately at transfer 01 and remaining until transfer 19 (Fig. 3). The communities exhibit a rebound, or recovery, in function after *E. coli* introduction was stopped at transfer 14 (comparing transfer 12 to 19, Fig. 3 C and E); at transfer 19 the PCA ellipses of metabolic profiles of the Recovery and the Control treatments overlap, while the Introduction treatment remains separated. The divide in community function between Control and Introduction treatments exists along principal component 1 (PC1) but not the second component (PC2) across transfers. However, the contribution of PC1 reduces overtime from 80% (transfer 1) to 53% (transfer 19) of the variation, while PC2 explains between 13% (transfer 1) to 31% (transfer 19); thus, strikingly, while for all transfers and treatments PC1 and PC2 explain more than 80% of the variation, the relative contribution of PC1 decreases and PC2 increases overtime. Moreover, the loadings on the metabolic compounds changed for PC1 from being all in the same direction at transfer 1 to contrasting directions at transfer 19, while the reverse was true for PC2, suggesting a change of metabolism over time.

**Figure 3:**
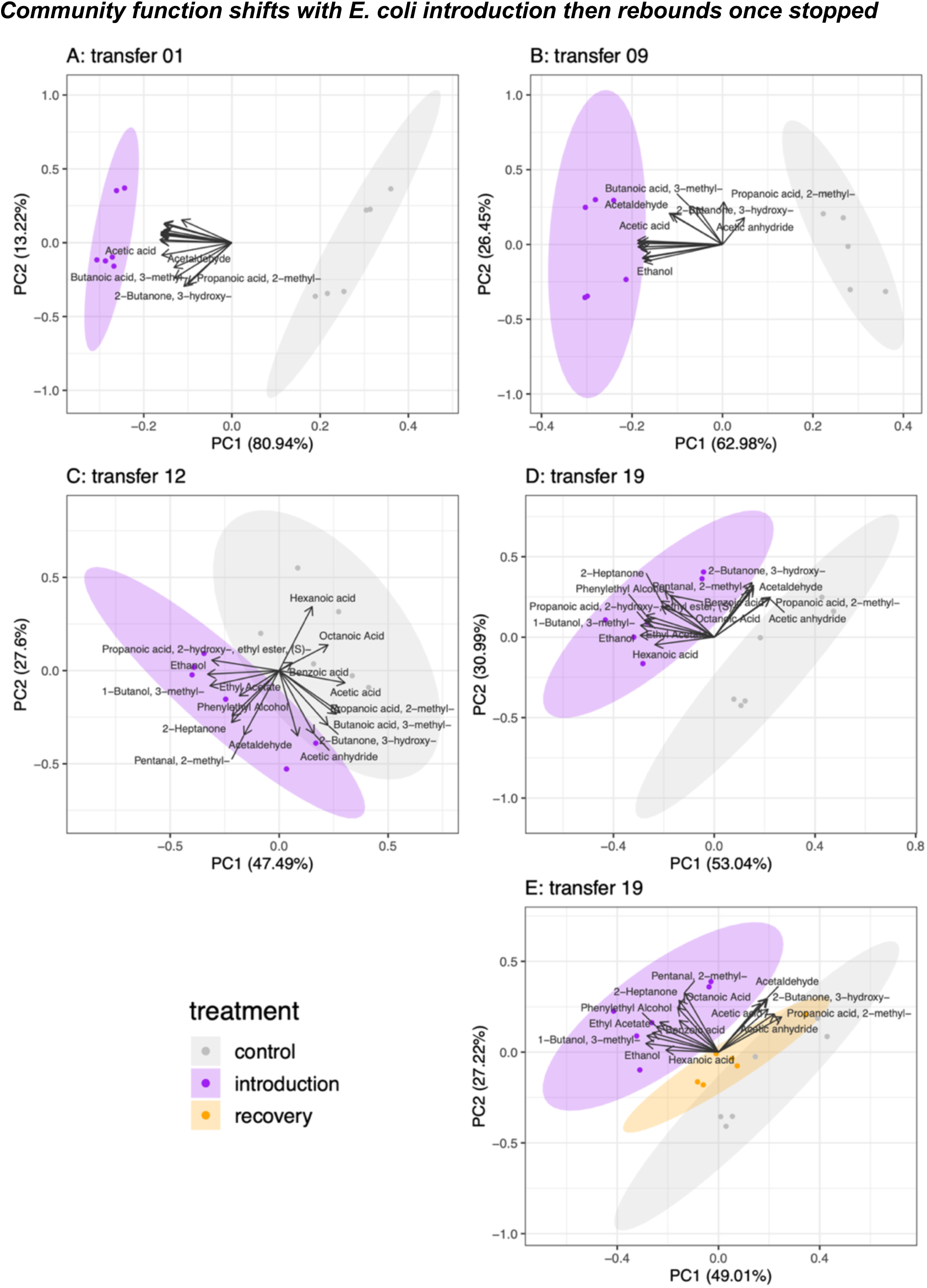
Community function is immediately impacted by E. coli introduction but rebounds following cessation. PCA analysis of metabolic profiles compiled from 16 volatile organic compounds (VOCs) using GC-MS analyses at transfer 1 (A), 9 (B), 12, (C) and 19 (D, E). Dots represent individual replicates. Ellipses are 95% confidence intervals. Direction of vectors indicate the compound’s contribution to either principal component, whereas the vector length indicates the amount of variation explained by the two plotted principal components. Some labels are not shown due to text overlap, but all compounds are listed in methods section. Transfer number refers to the end products of previous propagation (i.e., “transfer 9” are metabolic profiles of the end products from transfer 8). Details of compound loadings are found in supplementary Table S3.

Whether or not initially failed invaders may later establish in future introductions can be influenced by shifts in resource use of the resident community away (permissive) or towards overlapping (preventing) with the introduced species (Mallon *et al*. 2018). The same would apply for shifts in community function, which we measured as metabolic profiles of VOCs. We did not assess resource use in our experiment, and instead observed metabolic profiles to monitor changes in community function. The persistent division in metabolic profiles between Introduction *versus* Control or Recovery communities are characterised by a few VOCs which are repeatedly found at higher levels in the Introduction communities. These compounds include alcohols and esters potentially associated with Ehrlich pathway activity of the yeast *Galactomyces geotrichum* (ethanol, acetaldehyde, hexanoic acid-ethyl ester, 1-butanol 3-methyl) (Szudera-Kończal *et al*. 2020). It is possible that *E. coli* interacts with yeast and via changes in yeast abundances or metabolism, the community metabolic profiles are altered. Our previous work showed that the bacterial community compositions of Mabisi were not influenced by the presence-absence of *Galactomyces geotrichum* yeast (Leale *et al*. 2023); thus, as observed in this experiment, we would not expect to see shifts in *bacterial* community compositions even if yeast abundances were altered by *E. coli* introduction. Future work would provide insight on possible *E. coli* – yeast interactions in Mabisi.

We are unable to disentangle whether the observed shift and recovery in metabolic profiles are due to metabolism of resident community itself, or if the shifts are simply metabolic contributions of any transient *E. coli* growth. Sampling was done after 72 hours of fermentation, when the pH was already at minimal levels for approximately 24 hours (the largest pH drop in Mabisi fermentation occurs in first 48 hours (Groenenboom *et al*. 2020)). Although *E. coli* was not detected by full-length 16s rRNA gene sequencing at 72 hours (Fig. 3), it is possible that metabolism from growth earlier during fermentation remains evident in final metabolic profiles. Few VOCs can be linked to metabolic activity of *E. coli*. For instance, 2-heptanone has been found in *E. coli* cultures (Maddula *et al*. 2009), but this compound may as well originate from a lactic acid bacterium (Albright *et al*. 2020). The generality of metabolism of the species in our community in combination with increased intensity of most, or all compounds (i.e., Fig. 4 vectors similar in length and direction) means we cannot make specific associations between detected VOCs and certain community members such as *E. coli*. In hindsight, metabolic profiles of the Recovery treatment at transfer 15 would have shown whether community function immediately rebounded to similarity of Control lines, therefore lending evidence of *E. coli*’s metabolic contributions. Detecting the metabolic input of *E. coli* and yeast through transcriptomic data across transfers and within a cycle (i.e., 6, 12, 24 hours), could help us understand our observed metabolic plasticity.

**Figure 4:**
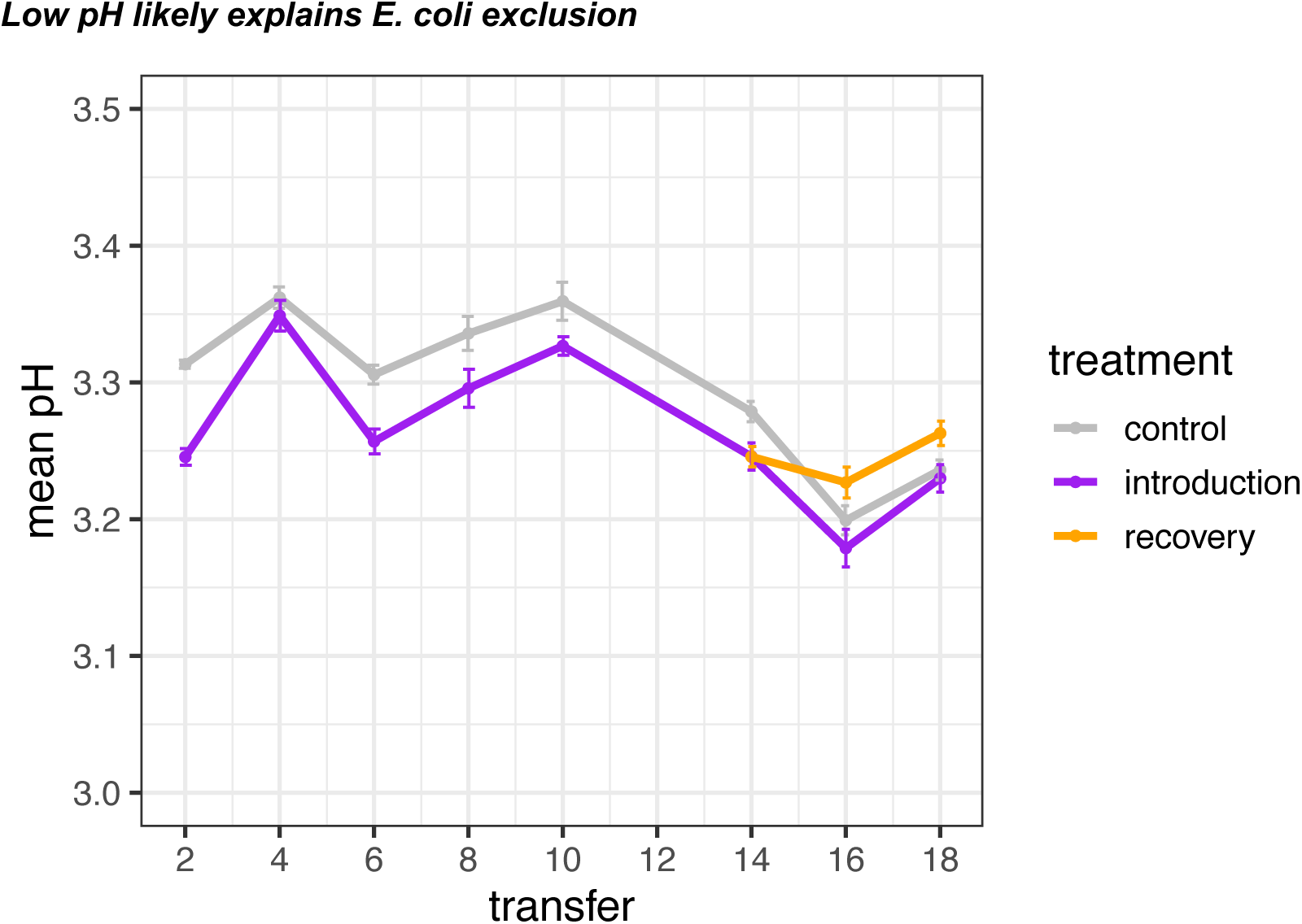
E. coli introduction slightly reduces pH. Measures of pH of Control, Introduction, and Recovery communities over 18 rounds of serial propagation. Points show mean value for the 6 replicate communities, vertical bars show standard error of the mean. Transfer number refers to the end products of previous propagation (i.e., pH “transfer 10” is the pH of end products from transfer 9).

We assessed pH over time, which is a higher order yet practical component of community function. A fast fermentation with a corresponding drop in pH is a contributing factor to the high biological safety of fermented food products (Mpofu *et al*. 2016) and consumer preference (Ott *et al*. 2000), including Mabisi (Sikombe *et al*. 2023). Low pH and high acidity are key determinants for the survival of *E. coli* in soil environments (Xing *et al*. 2021). In our experiment, the fully grown Mabisi culture had a pH of around 3.25 (Fig. 4) which is well below the optimal pH of *E. coli* of 6.5 to 7.5 (Davey 1994), lending explanation to the inability of *E. coli* to establish (Fig. 3). Interestingly, the Introduction treatment had a significant lower pH as compared to the Control treatment (Table S4: ANOVA excluding Recovery treatment, effect of treatment (p < 0.0001, F = 23.51, DF = 1), transfer (p < 0.0001, F = 34.40, DF = 7), and treatment x transfer interaction (p = 0.06052, F = 2.04, DF = 7)), with the exception at transfer 4 and until later time points when Control and Introduction treatments were similar (Table S5: Tukey’s pairwise comparison Control versus Introduction p > 0.05 at transfers 4, 16, and 18). *E. coli* does grow alone in milk and drops the pH because of metabolic products (Fig. S2 and S3), so the small drop in pH in the Introduction treatment is likely simply the result of an overall higher number of cells (i.e., CFU Mabisi community + CFU *E. coli*), and not any interaction of the resident community with *E. coli.* We consider no biological significance in interpreting such a minor reduction of pH (< 0.05 pH units) resulting from *E. coli* introduction.

## GENERAL DISCUSSION

The main aim of this study was to investigate repeated introduction of a novel species to a microbial community over many generations. We were interested in whether a novel species not predicted to establish may still alter the resident community composition and function over time when repeatedly introduced, and if there would be recovery once introduction of the novel species was stopped. We immediately observed alteration of community function (i.e., metabolic profiles) followed by recovery once *E. coli* introduction ceased. The observed shifts in community function in response to novel species introduction was not similarly reflected in bacterial community species composition. No change in the bacterial community composition was detected; however, the likelihood of assessing possible community composition shifts was reduced due to unexpectedly low starting species diversity in our ancestor community. Overall, we observed a shift and rebound of metabolic profiles when *E. coli* was introduced without changes in species composition.

We cannot yet realise the several potential existing mechanisms by which the shift and rebound in community function occurs. Our first explanation of the observed metabolic shift is an interaction between *E. coli* and *G. geotrichum* yeast found in our Mabisi samples (Leale et al. 2023). We secondly speculate that the metabolism of the resident community was changed by *E. coli*’s presence, yet the species compositions themselves remained unaltered. Thirdly, there may be changes at the genotypic level which are influencing community function, but they were not detected in our analysis level of full-length 16s rRNA gene sequencing. The low diversity we found, and its consequent limitations, emphasises the importance of the choice of resident community and novel introduced species in future research. Next steps to investigate these three possible mechanisms could involve a multitude of approaches; first estimating yeast abundances through selective agar plating, meta-transcriptomics could then further provide detailed insight into the metabolic contributions of community members, including *E. coli* and yeast, and lastly, metagenomics could show hidden diversity and dynamics of bacterial sub-species genotypes not detected here. Microorganisms can change their metabolism in response to their biotic environment and community member interactions (van Rijswijck *et al*. 2017). To investigate possible “metabolic plasticity”– a cell’s ability to change their nutrient conversion (Fendt, Frezza and Erez 2020) – to *E. coli* introduction, we should also demonstrate stability in the community genotypes. There are many exciting future directions, yet what we can say for now from our experiment is that *E. coli* introduction produced community level metabolic changes without sufficient pressure to alter bacterial species sorting.

A community’s resistance against novel species may be strengthened or weakened with repeated exposure overtime. Previous research on unsuccessful invasions (Mallon *et al*. 2018; Amor, Ratzke and Gore 2020; Xing *et al*. 2021) brings to question whether potential compositional and functional shifts in the resident community are beneficial to the resident community, or to the introduced species? We pose three hypotheses; either the shift makes the novel species more successful at establishing later, the reverse – that the resident community is now more stable against future introductions, or thirdly the shift could have no effect. The resident community may adapt ecologically and genetically in response to selection by continued introduction of a novel species over many generations. It is suggested that evolutionary processes can contribute to stability, such as to invasion, at the community level (Jousset *et al*. 2011; Kinnison, Hairston Jr and Hendry 2015; Hendry 2019; Vila *et al*. 2019). Specialization, generalization, and diversification are evolutionary trajectories that can create complementarity in resource use or overall niche space and enhance exclusion of a novel species (Gravel *et al*. 2011; van Moorsel *et al*. 2021). Simply ecological processes of species sorting, without genotypic changes, could also improve total resource use and leave less available niche space through similar processes of complementarity. If true, we would predict that communities initially unable to resist establishment by an introduced species will increase resistance after experimental evolution due to reducing remaining resources. Our results do not support either prediction about the fate of establishment overtime by an introduced species – *E. coli* establishment was neither lower nor greater after 19 propagation cycles (approximately 110 generations). In general, introduced species seldomly persist, or at least do not induce lasting changes to community function. It is proposed that biotic interactions between the introduced species and resident community are more important for invader persistence than the dosage pressure of introduction (Albright *et al*. 2020, 2022). Our results support such a system-specific fitness perspective, as the fitness of *E. coli* in the Mabisi environment appears to be too low even with its high initial inoculation levels (approximately 50% *E. coli* CFU/mL).

Controlled laboratory microbial evolution experiments tracking establishment overtime of a repeatedly introduced novel species could identify critical tipping points in resistance of the resident community. Our experiments with an *E. coli* invader in the highly acidic Mabisi environment did not result in long-term establishment of an initially unsuccessful invader. However, an exciting future avenue to pursue is using an introduced species that is ecologically alike to, but not found in, our Mabisi community, such as *a Lactococcus*. Introducing *Lactococcus* is also relevant since natural Mabisi communities contain either dominant levels of lactococci or lactobacilli, or they coexist at similar abundances (Schoustra *et al*. 2013; Moonga *et al*. 2021). It is not yet entirely clear what selection factors determine the dominance of lactobacilli or lactococci in a Mabisi community, whether frequency dependence contributes, if it is a stochastic drift process, or if temperature is a main factor (Moonga *et al*. 2021). We could begin investigating their coexistence in Mabisi by following the fate of an introduced *Lactococcus* strain in our communities where it is not initially present (Fig. 2 this paper and Fig. 2 (Leale *et al*. 2023)) under varied temperatures. Similar work traced the fate of an introduced probiotic lactic acid bacteria in a traditional fermented milk from Senegal. The probiotic strain was detected at minimal levels (<1%) after only the first fermentation cycle, then was quickly lost following the second back-slop propagation (Groenenboom *et al*. 2019). The remaining question then is if stable establishment would eventually result from repeated introduction of the probiotic strain at every fermentation cycle?

Microbial communities play critical roles to environmental, industrial, and human ecosystem functions, all of which face disturbance by species introductions (Kinnunen *et al*. 2016; Bolius, Wiedner and Weithoff 2019; Gu *et al*. 2019; Libertucci and Young 2019; Oro *et al*. 2019; Albright *et al*. 2020). An introduced species may not successfully establish yet still impact the dynamics of species interactions and have a potentially lasting influence on the overall community composition and function (Mallon *et al*. 2018; Amor, Ratzke and Gore 2020; Xing *et al*. 2021). Community composition and function may not respond in parallel to an introduced species, therefore the engineering of new microbial ecosystems or the preservation of existing ones requires concurrently investigating both components. Ours and others’ observed change in function yet not composition in response to disturbance (Sjöstedt *et al*. 2018) is an important consideration for industrial applications; for example, a bioreactor or food product may not be detectably contaminated by an unwanted organism, yet the function of interest is still impacted. There is additionally a critical emphasis to expand the time scales of measuring microbial community responses to perturbations and testing the effect of multiple, rather than single incidents (Vila *et al*. 2019; Philippot, Griffiths and Langenheder 2021; Xing *et al*. 2021; Albright *et al*. 2022). Soil systems heavily represent existing research on microbial community resistance to invasion (van Elsas *et al*. 2007; Allison and Martiny 2008; Shade *et al*. 2012; Yao *et al*. 2014; Mallon *et al*. 2018; Amor, Ratzke and Gore 2020; Xing *et al*. 2021), thus our work expands upon the field by using a unique fermented food model system. Our study’s effort to measure microbial community composition and function over a hundred generations, including following cessation of repeated novel species introduction, contributes to understanding long-term community stability to sustained disturbances.

## METHODS

### Preparation for transfer 0

A fermented Mabisi sample stored at –80 C in 50%v/v glycerol was defrosted and 800uL inoculated into 40mL of UHT full fat milk (Campina Langlekker volle melk) then incubated unshaken at 28°C for 72 hours. On the same day, a glycerol stock of *E. coli* (strain DSM498) was inoculated into 2mL of BHI broth (VWR 84626.0500) and grown for 72 hours shaken at 37°C. The choice of 72 hours for *E. coli* growth culture was done instead of 24 or 48 hours because of feasibility with laboratory working hours and weekends. *E. coli* cultures were therefore always at stationary phase when inoculating into milk with Mabisi. Cell density of *E. coli* after 72-hours in BHI broth was estimated by plate counts to be 3.55 x 10^9^ CFU/mL (Fig. S3).

### Transfer 0 inoculations

From the 72-hour fermented Mabisi culture and *E. coli* culture, two treatments were created with six replicate lines each. Following inversion and mixing of the Mabisi culture, 400 uL of culture was added to 40 mL of fresh full milk for both the Control (CON) and Introduction (INT) replicates. The Introduction treatment additionally received 40 uL of *E. coli* culture, producing an estimated 3.55 x 10^6^ CFU/mL of *E. coli* at Transfer 0 in 40 mL of milk; this concentration is greater than field study estimates of enterobacteria found in raw milk used for Mabisi production in Zambia (Schoustra *et al*. 2022). Samples then underwent repeated 72-hour fermentation cycles at 28 °C for 18 transfers (approximately 110 generations); fermented cultures were mixed then 400 uL transferred to 40 mL of new milk, with 40 uL of new 72-hour *E. coli* culture also added to Introduction treatment replicates at every transfer. For logistical purposes, after every second transfer final cultures were inoculated into fresh milk and stored at 4 °C for 24 hours before being moved to 28 °C. Therefore, a “true” cycle was completed after every second transfer (i.e., 7 days). The 72-hour *E. coli* culture was repeatedly grown fresh from the same glycerol stock for each inoculation in the Introduction treatment. At transfer 14, cultures from each replicate of the Introduction treatment were used to create a third “Recovery” treatment, where fresh *E. coli* culture was not added at the remaining transfers. Fresh samples of final products were archived at transfers 1, 5, 9, 12, and 19, with the following archived: with glycerol at –80 °C (1.27 mL culture + 0.63 mL 85% glycerol), and without glycerol at –20 °C for DNA analysis and GC-MS analysis (remaining volume in tube). Please note that archive labelling was done as follows: t9 = final product from transfer 8 (i.e., the Mabisi culture used to inoculate at transfer 9), t19 = final product from transfer 18. The pH was also measured at every second transfer.

### GC-MS Analysis of VOCs

Larger tubes of samples without glycerol at –20 °C were defrosted at 4 °C, thoroughly mixed, 1.8 mL pipetted into headspace vials, then stored at –20 °C until analysis. A sample of 1.8 mL Jumbo Brand Volle Melk and 1.8 mL Jumbo Kefir Naturel were included with every time point as controls. After incubating for 20 minutes at 60 °C, a SPME fibre (Car/DVB/PDMS, Suppelco) extracted volatiles for 20 minutes at 60 °C. Volatiles were desorbed from the fibre under the following conditions: Stabilwax-DA-Crossbond-Carbowax-polyethylene-glycol column (2 min), PTV split mode at a ratio of 1:25 (heated to 250 °C), helium carrier gas at 1.2 mL/min, GC over temperature at 35 °C (2 min) raised to 240 °C (10 C/min), kept at 240 °C (5min). Mass spectral data was collected over a range of 33-250 m/z in full scan mode with 3.0030 scans/seconds. Results were analysed with Chromeleon 7.2 CDS Software (ThermoFisher) where the following signal peaks were identified as volatile metabolites according to their elution time and mass spectral data: acetaldehyde; Ethyl Acetate; Ethanol; 2-Heptanone; 1-Butanol, 3-methyl-; 2-Butanone, 3-hydroxy-; Propanoic acid, 2-hydroxy-, ethyl ester, (S)-; Acetic anhydride; Pentanal, 2-methyl-; Acetic acid; Propanoic acid, 2-methyl-; Butanoic acid, 3-methyl-; Hexanoic acid; Phenylethyl Alcohol; Octanoic Acid; Benzoic acid. MS quantification peak counts were exported to Excel. Data was first normalised by compound using the calculation 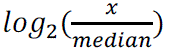, where x is the quantification peak count for a given compound in each sample (i.e., area under compound peak), and the median ion count across all samples for that compound.

### DNA extraction

Larger tubes of samples without glycerol at –20 °C were defrosted at 4°C, thoroughly mixed, and ∼1.75 mL pipetted into centrifuge tube and stored at –20 C. On the day of extraction, the sample was defrosted and spun down (2 min, 13000 g), then the supernatant and curd removed with a sterile scoopula. Cells were re-suspended in a mix of 64 μL EDTA (0.5 M, pH 8), 160 μL Nucleic Lysis Solution, 5 μL RNAse, 120 μL lysozyme [10 mg/mL] and 40 μL pronase E [10 mg/mL] and incubated for 60 minutes at 37 °C with agitation of 350 rpm. Cells were dislodged by manually flicking the tube occasionally during incubation, to improve mixing with suspension mixture. Bead beading (MP Biomedical FastPrep-25 5G device) was then performed for 3 minutes (1 minute, 5-minutes rest, repeated three times) with sand sized beads, then 400 μL ice-cold ammonium acetate (5 M) added, and the mixture immediately cooled on ice for 15 minutes. The mixture was spun down (4 min, 13000 g) and 650 μL transferred to a 2 mL flat bottom 96-well plate (Greiner #789271). The plate was sealed tightly with a rubber cover mat, inverted to mix, incubated for 10min on ice, then centrifuged (5 min, 3000 g). The six replicates per treatment were split so that there were three replicates per 96 well plate; this organisation remained for all future steps, including Oxford Nanopore sequencing.

In a new 1 mL 96-well round bottom plate (Greiner #780201), 400 µL of supernatant from the previous plate and 400 µL of homemade SPRI beads were added (1 mL Sera-Mag SpeedBeads (Cytiva, Marlborough, MA, USA) cleaned and dissolved in 50mL end volume containing 2.5 M NaCl, 20 mM PEG, 10mM Tris-HCl and 1mM EDTA). The plate was sealed tightly with a rubber cover mat, placed on its side and shaken at 50 rpm for 1 hour. The plate was then left on the 96 well magnet (Thermo Fisher #AM10027) for 5-10 minutes, and the supernatant was poured off (plate inverted at sink while held on magnet). A wash was performed three times by adding each time 500 µL of 80% ethanol, sealing the plate tightly with the rubber cover mat, vortexing, returning plate to magnet, then pouring off ethanol. After the third rinse, beads were left to dry for 10-30 min and 50 µL of sterile milliQ water added. The plate was shaken horizontally to resuspend the beads, left for 10min at room temperature, then returned to the magnet for 5-15min. While remaining on the magnet, 45 µL of the eluate containing genomic DNA was transferred to a regular 96 well plate and stored at –20 °C. DNA concentrations were then measured by Invitrogen Quant-iT PicoGreen according to manufacturer instructions (ThermoFisher #P7589).

### Nanopore MinIon protocol

This protocol was adapted from Beekman et al. 2022 and Leale et al. 2023 (Beekman *et al*. 2022; Leale *et al*. 2023). DNA concentrations in these steps, with the exception of the final pooled sample, were measured by Invitrogen Quant-iT PicoGreen.

#### Tailed PCR reaction

The first step in Nanopore sequencing was a PCR reaction of 19 cycles using Nanopore specific tailed primers. Three positive controls were included – the ZymoBIOMICS Microbial Community DNA Standard D6305, a previously sequenced Mabisi sample (community “F1.1” from Leale et al. 2024) (Leale *et al*. 2023), and a personally created sample titled “M+”. The M+ control contained cleaned colony PCR products of the full 16s rRNA gene (27F, 1492R primers) of five species, combined in equal proportions by measured DNA concentration – *Limosilactobacillus fermentum* (DSM20052), *Lactobacillus helveticus* (DSM20075), *Lactobacillus delbrueckii* (DSM20072), *Lactococcus lactis* (DSM20481), *Acetobacter orientalis* (DSM15550), *Acetobacter lovaniensis* (DSM4491). The tailed primer PCR reaction was as follows:

##### Primers

Nanopore tailed forward: 5’ TTTCTGTTGGTGCTGATATTGC-[27F] 3’

Nanopore tailed reverse: 5’ ACTTGCCTGTCGCTCTATCTTC-[1492R] 3’

27F: 5’ AGA GTT TGA TCC TGG CTC AG 3’

1492R: 5’ TAC GGY TAC CTT GTT ACG ACT T 3’

##### Tailed reaction reagents

1 µL – DNA [0.5ng/µL]

12.5 µL – Phusion High Fidelity PCR 2X master mix (ThermoFisher)

1.25 µL – forward tailed primer [10uM]

1.25 µL – reverse tailed primer [10uM]

9 µL – MilliQ water

##### Tailed cycle conditions

98 °C 10 sec

98 °C 5 sec (19X)

57 °C 5 sec (19X)

72 °C 30 sec (19X)

72 °C 1 min

12 °C infinity

The tailed PCR reaction was performed another two times, resulting in three separate tailed primer PCR products per sample. Placement of PCR tubes in machine was adjusted for each PCR to avoid edge effects. Each amplified DNA sample was all visualized on 1% agarose gel to confirm successful amplification, then 8 µL of each PCR reaction were combined. A total of 24 µL of amplified DNA per samples was used for the clean-up.

#### PCR Clean-up

For each sample, the 24 µL of amplified DNA was cleaned with 24 µL of homemade SPRI beads (i.e., 1:1 ratio) (1 ml Sera-Mag SpeedBeads (Cytiva, Marlborough, MA, USA) cleaned and dissolved in 50 ml end volume containing 2.5 M NaCl 20 mM PEG, 10mM Tris-HCl and 1 mM EDTA) and eluted into 20 µL of MilliQ water. DNA concentration of cleaned amplicons was measured using Invitrogen Quant-iT PicoGreen. A new dilution of 15 µL of the cleaned, amplified 16S PCR product was made into a new diluted sample with DNA concentration 0.5 nM.

#### Barcoding

The PCR for each sample was barcoded to enable pooling using the PCR Barcoding Expansion 1-96 Kit (Oxford Nanopore Technologies). Reaction volumes were adapted from the Nanopore barcoding protocol to save in reagents used. The reaction was as follows with a unique barcode per sample:

##### Barcoding PCR (per reaction/sample)

0.3 µL barcode (Oxford Nanopore Technologies)

7.2 µL [0.5nM] cleaned PCR

7.5 µL LongAmp Taq 2x Master Mix (New England Biolabs)

##### Cycle conditions

95 °C – 3 min (x1)

95 °C – 15 sec (x16)

62 °C – 15 sec (x16)

65 °C – 1.5 min (x16)

65 °C – 2 min (x1) 4 °C – infinity

Before pooling of the PCR barcoded products, each was visualised on 1% agarose gel. All samples were combined with 2 µL, apart from samples with fainter bands that were subjectively determined to require 3 µL (1-F1, 1-S1, 5-L1), 4 µL (5-S1, 1-M5), or 5 µL (1-S2, 1-F5, negative control). The pooled sample (total volume = 208 µL) was cleaned using homemade SPRI beads in 1:1 volumetric ratio and eluted in 200 µL of milliQ water. The DNA concentration of the cleaned, pooled sample was measured on the Qubit 2.0 fluorometer using dsDNA High Sensitivity Assay kit (Thermo Fisher).

In 47 µL of milliQ water, 1 ug of the barcoded, pooled, cleaned library was prepared. From here, the library was repaired, end-prepped and adaptor ligated according to the Oxford Nanopore Technologies ligation sequencing kit V14 (SQK-LSK114), following the protocol: ligation-sequencing-amplicons-sqk-lsk114-ACDE_9163_v114_revJ_29Jun2022-minion”. Reagents used were NEBNext FFPE DNA Repair Buffer (E7181A), NEBNext FFPE DNA End Repair Mix (E7182A), NEBNext Ultra II End Prep Reaction Buffer (E7183A), and NEBNext Ultra II End Prep Enzyme Mix (E7184A) (New England Biolabs).

The prepared library was loaded on a R10.4.01 SpotON Flow Cell (FLO-MIN114) on a MinION MK1c sequencing device (Oxford Nanopore Technologies) at 5 kHz sampling rate and 400 bases/second reading rate. Base-calling was performed shortly after using the “fast base-calling” setting.

Barcodes removed from final analyses were plate 1 and 2, bc 93 = “F1.1” natural Mabisi community DNA (Leale *et al*. 2023), bc94 = “M+” mock synthetic Mabisi community created from DNA of DSMZ strains, bc95 = ZymoBIOMICS Microbial Community DNA Standard, bc96 = negative control.

#### Bioinformatics

To assign 16S rRNA sequences to taxonomic identity, we first downloaded the SILVA reference database (Quast *et al*. 2013). Since we were interested in variation above the species level, a custom database was produced using vsearch to cluster 16S rRNA sequences at a 95% similarity threshold (Rognes *et al*. 2016). Reads were aligned to this database using minimap2 (Li 2018) and the best matching cluster was assigned as the taxonomic identity for each read. Details of sequence compositions for main clusters are found in Table S3 from Leale et al. 2023 (Leale *et al*. 2023).

Clusters to which less than five reads matched were assumed to be trace contamination and discarded. The overall frequency of clusters with <1% abundance is similar across diversity levels, as evidenced by the amount of “missing reads” to complete 1.00 abundance on bar plot (i.e., blank space at top of bar plot).

Due to a complication between the newer MinIon flowcell and older machine versions, the base-calling and then bioinformatic analysis for the first 96-well plate posed substantial complications. This was however solved for the second batch of samples (replicates 4, 5, 6). Since there was minimal species diversity in, nor variation between, replicate populations 4, 5, and 6, we decided to only consider these three. The presence of only two lactic acid bacterium types in the in our starting Transfer 0 community was not expected, which we yet to have a clear explanation for. Our DNA extraction method differed from Leale et al. 2024, but a preliminary test confirmed that the abundances of types did not significantly differ between extraction techniques, reassuring our approach.

The yeast present in the Mabisi communities used in this study was isolated and identified as *Geotrichum candidum* using amplification and sequencing of the ITS (internal transcribed spacer) region (top match GenBank: MK967716.1). Yeast was visibly seen growing at the air interface of both Control and Introduction treatments.

#### Statistical analyses

All analyses were performed in R version 4.3.2 (R Core Team 2023). Significance was defined for all analyses as p-value < 0.05. Results outputs and details of statistical analysis are found in Supplementary Material.

#### Community compositions

A PERMANOVA was performed on the full data set of community compositions to evaluate the effect of treatment, transfer, and their interaction on community composition. The “adonis” function from “vegan” R package was used, with “bray” method. Community richness was also calculated using the “vegan” package’s “specnumber” function (Oksanen *et al*. 2022).

#### Acidity

Excluding Recovery treatment populations, a two-way ANOVA of community*transfer was performed on pH values using the “lm” function from “lme4” R package, type 3 partial sum of squares (Bates *et al*. 2015). Post-hoc Tukey’s pairwise comparisons with adjusted p-values were then performed using the “emmeans” function from “vegan” R package at 95% confidence intervals (Oksanen *et al*. 2022).

## AUTHOR CONTRIBUTIONS

Experimental design and conceptual ideas were developed by AML and SS. Experiments and data collection were performed by AML and FRM. Data analysis was completed by AML. Manuscript written and edited by AML, SS, EJS, BZ.

## ACKNOWLEDGMENTS

A big thanks to Dr. Ben Auxier for creating the bioinformatic analysis pipeline and his support with data analysis and visualisation. We additionally thank Judith Wolkers-Rooijackers for running the GC-MS machine.

## DATA AVAILABILITY

Source data files and codes available on GitHub (amleale/ecoli_mabisi_2022): https://github.com/amleale/ecoli_mabisi_2022. Raw amplicon sequence data for the communities are available under NCBI BioProject PRJNA1096163 with each barcode available separately under SAMN40628846 – SAMN40629034.

## SUPPLEMENTARY

### Statistical Analyses – supplementary details

#### 1. COMMUNITY COMPOSITIONS

*“COMMUNITY” refers to the treatment of “Control”, “Introduction”, and “Recovery”*.

*adonis2(wide[,9:12] ∼ COMMUNITY*TRANSFER, data = wide[,2:3], method =*

*“bray”)*

*Permutation test for adonis under reduced model*

*Terms added sequentially (first to last)*

*Permutation: free*

*Number of permutations: 999*

**Table S1:**
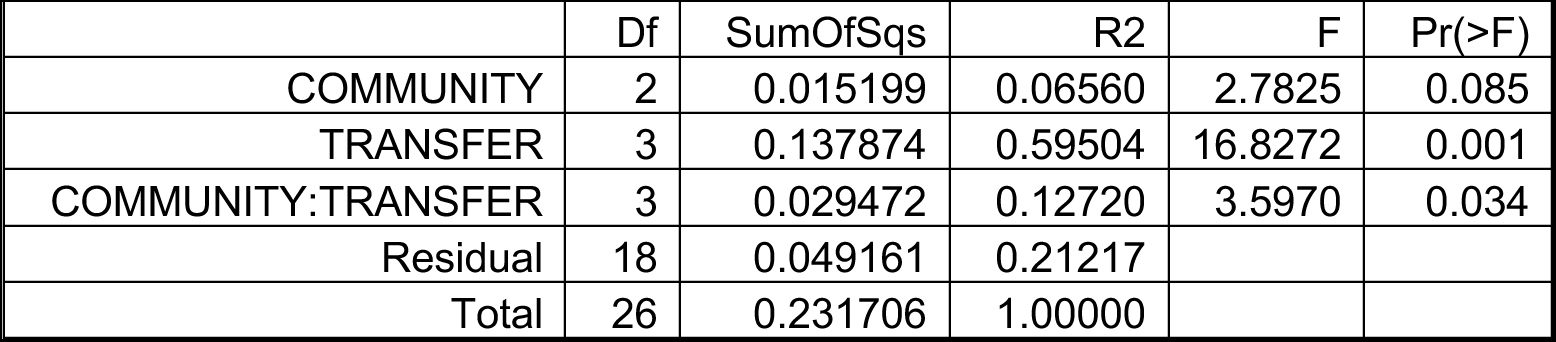
Output of PERMANOVA tests of full 16s rRNA gene community compositions.

|  | Df | SumOfSqs | R2 | F | Pr(>F) |
| --- | --- | --- | --- | --- | --- |
| COMMUNITY | 2 | 0.015199 | 0.06560 | 2.7825 | 0.085 |
| TRANSFER | 3 | 0.137874 | 0.59504 | 16.8272 | 0.001 |
| COMMUNITY:TRANSFER | 3 | 0.029472 | 0.12720 | 3.5970 | 0.034 |
| Residual | 18 | 0.049161 | 0.21217 |  |  |
| Total | 26 | 0.231706 | 1.00000 |  |  |

**Table S2:** Mean species richness from full 16s rRNA gene community compositions.

|  | Control | Invasion | Recovery |
| --- | --- | --- | --- |
| t01 | 2.67 | 3 | n/a |
| t09 | 2.33 | 2 | n/a |
| t12 | 1.67 | 1.67 | n/a |
| t19 | 1.33 | 1.67 | 2.33 |

#### 2. METABOLIC PROFILES

**Figure S1:**
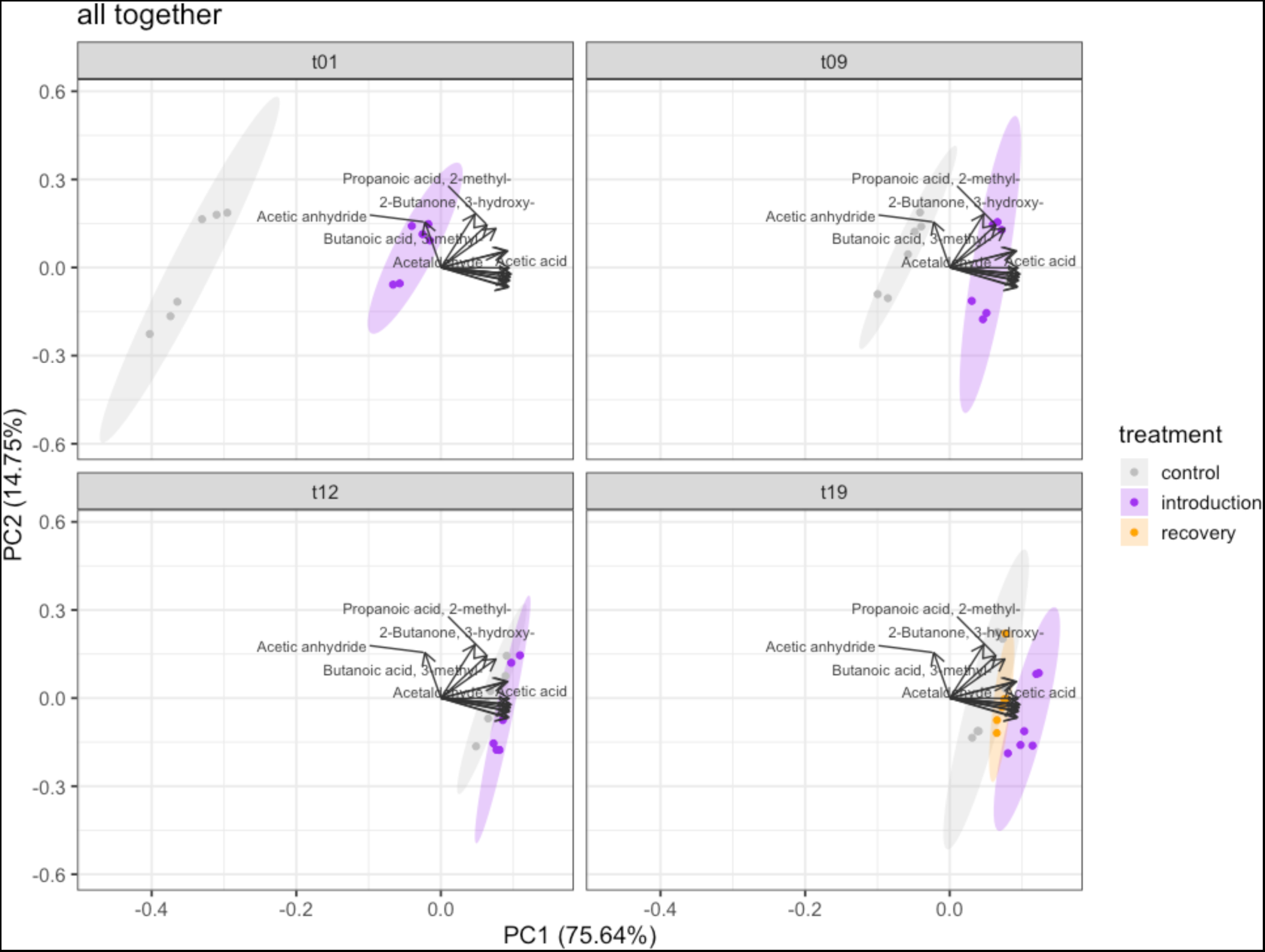
PCA of metabolic profiles calculated for all data points combined and plotted per transfer, rather than a separate PCA analysed for each transfer. T01 = transfer 1, t09 = transfer 9, t12 = transfer 12, t19 = transfer 19.

**Table S3:** PCA loadings of aroma compounds in figure 3.

|  | transfer 1 (fig3.a) |  |  | transfer 9 (fig3.b) |  |  | transfer 12 (fig3.c) |  |  | transfer 19 (fig3d) |  |  |
| --- | --- | --- | --- | --- | --- | --- | --- | --- | --- | --- | --- | --- |
|  | PC1 | PC2 | PC3 | PC1 | PC2 | PC3 | PC1 | PC2 | PC3 | PC1 | PC2 | PC3 |
| Acetaldehyde | -0,22 | -0,29 | -0,06 | -0,19 | 0,34 | 0,22 | 0,09 | -0,38 | 0,20 | 0,17 | 0,36 | -0,01 |
| Ethyl Acetate | -0,26 | 0,15 | -0,19 | -0,29 | -0,14 | -0,02 | -0,20 | -0,15 | 0,31 | -0,30 | 0,11 | -0,24 |
| Ethanol | -0,26 | 0,23 | -0,15 | -0,29 | -0,19 | -0,04 | -0,36 | -0,02 | 0,13 | -0,33 | 0,01 | -0,16 |
| 2-Heptanone | -0,27 | 0,10 | -0,07 | -0,31 | -0,03 | -0,09 | -0,24 | -0,30 | 0,12 | -0,24 | 0,25 | 0,07 |
| 1-Butanol, 3-methyl- | -0,26 | 0,21 | -0,11 | -0,29 | -0,15 | 0,09 | -0,34 | -0,09 | -0,03 | -0,33 | 0,08 | -0,22 |
| 2-Butanone, 3-hydroxy- | -0,18 | -0,50 | -0,01 | -0,09 | 0,41 | 0,41 | 0,24 | -0,32 | -0,08 | 0,17 | 0,38 | -0,02 |
| Propanoic acid, 2-hydroxy-, ethyl ester, (S)- | -0,27 | 0,12 | 0,00 | -0,29 | -0,15 | -0,02 | -0,33 | 0,06 | 0,15 | -0,31 | 0,12 | -0,02 |
| Acetic anhydride | -0,19 | 0,26 | 0,91 | 0,08 | 0,30 | -0,73 | 0,17 | -0,37 | 0,14 | 0,25 | 0,27 | -0,06 |
| Pentanal, 2-methyl- | -0,28 | 0,03 | -0,02 | -0,31 | 0,04 | -0,09 | -0,17 | -0,37 | 0,19 | -0,22 | 0,32 | -0,01 |
| Acetic acid | -0,27 | -0,14 | -0,08 | -0,18 | 0,33 | -0,23 | 0,33 | -0,07 | 0,14 | 0,16 | 0,32 | 0,41 |
| Propanoic acid, 2-methyl- | -0,17 | -0,48 | 0,26 | 0,00 | 0,47 | -0,06 | 0,30 | -0,24 | -0,09 | 0,25 | 0,28 | -0,05 |
| Butanoic acid, 3-methyl- | -0,22 | -0,41 | 0,08 | -0,12 | 0,43 | 0,21 | 0,28 | -0,25 | -0,13 | 0,18 | 0,34 | -0,23 |
| Hexanoic acid | -0,27 | 0,11 | -0,05 | -0,30 | -0,03 | -0,21 | 0,16 | 0,37 | 0,29 | -0,26 | -0,05 | 0,57 |
| Phenylethyl Alcohol | -0,27 | 0,12 | -0,07 | -0,30 | -0,06 | 0,17 | -0,23 | -0,26 | 0,01 | -0,30 | 0,15 | -0,26 |
| Octanoic Acid | -0,27 | 0,09 | -0,06 | -0,30 | 0,03 | -0,21 | 0,24 | 0,15 | 0,47 | -0,24 | 0,21 | 0,49 |
| Benzoic acid | -0,27 | 0,02 | -0,02 | -0,31 | 0,01 | 0,02 | 0,06 | 0,05 | 0,63 | -0,19 | 0,29 | -0,04 |

#### 3. ACIDITY

Recovery treatment excluded.

lm_ph <-lm(ph ∼ treatment*transfer, data = invade)

> Anova(lm_ph, type = 3) Anova Table (Type III tests)

emmeans(lm_ph, pairwise ∼ treatment|transfer)

**Table S4:** Output of ANOVA analysis for measured pH values.

|  | SumSq | Df | F value | Pr(>F) |
| --- | --- | --- | --- | --- |
| (Intercept) | 65.869 | 1 | 1.1219e+05 | < 2.2e-16 |
| treatment | 0.014 | 1 | 2.3512e+01 | 6.004e-06 |
| transfer | 0.141 | 7 | 3.4402e+01 | < 2.2e-16 |
| treatment:transfer | 0.008 | 7 | 2.0358e+00 | 0.06052 |
| Residuals | 0.047 | 80 |  |  |

**Table S5:** Output of Tukey’s pairwise comparisons between communities for measured pH values.

| transfer | contrast | estimate | SE | df | t.ratio | p.value |
| --- | --- | --- | --- | --- | --- | --- |
| t02 | con-inv | 0.06783 | 0.014 | 80 | 4.849 | <.0001 |
| t04 | con-inv | 0.01317 | 0.014 | 243 | 0.941 | 0.3494 |
| t06 | con-inv | 0.04883 | 0.014 | 80 | 3.491 | 0.0008 |
| t08 | con-inv | 0.04017 | 0.014 | 80 | 2.871 | 0.0052 |
| t10 | con-inv | 0.03267 | 0.014 | 80 | 2.335 | 0.0220 |
| t14 | con-inv | 0.03283 | 0.014 | 80 | 2.347 | 0.0214 |
| t16 | con-inv | 0.02033 | 0.014 | 80 | 1.453 | 0.1500 |
| t18 | con-inv | 0.00617 | 0.014 | 80 | 0.441 | 0.6605 |

#### 4. E.COLI ALONE IN MILK

**Figure S2:**
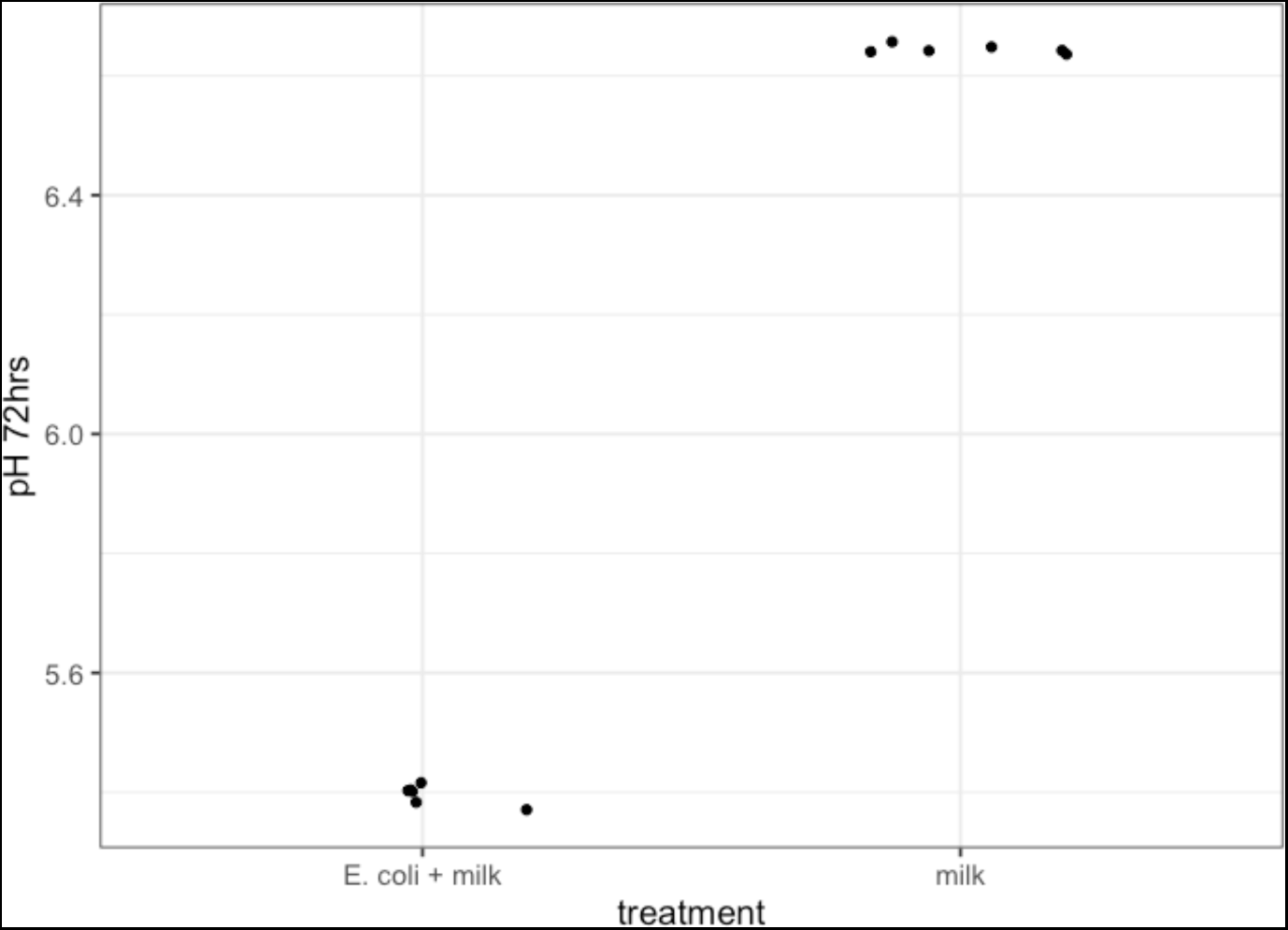
pH of milk inoculated or not with *E. coli* after 72 hours of fermentation at 28°C. Replication = 6, p-value < 2.2e-16.

**Figure S3:**
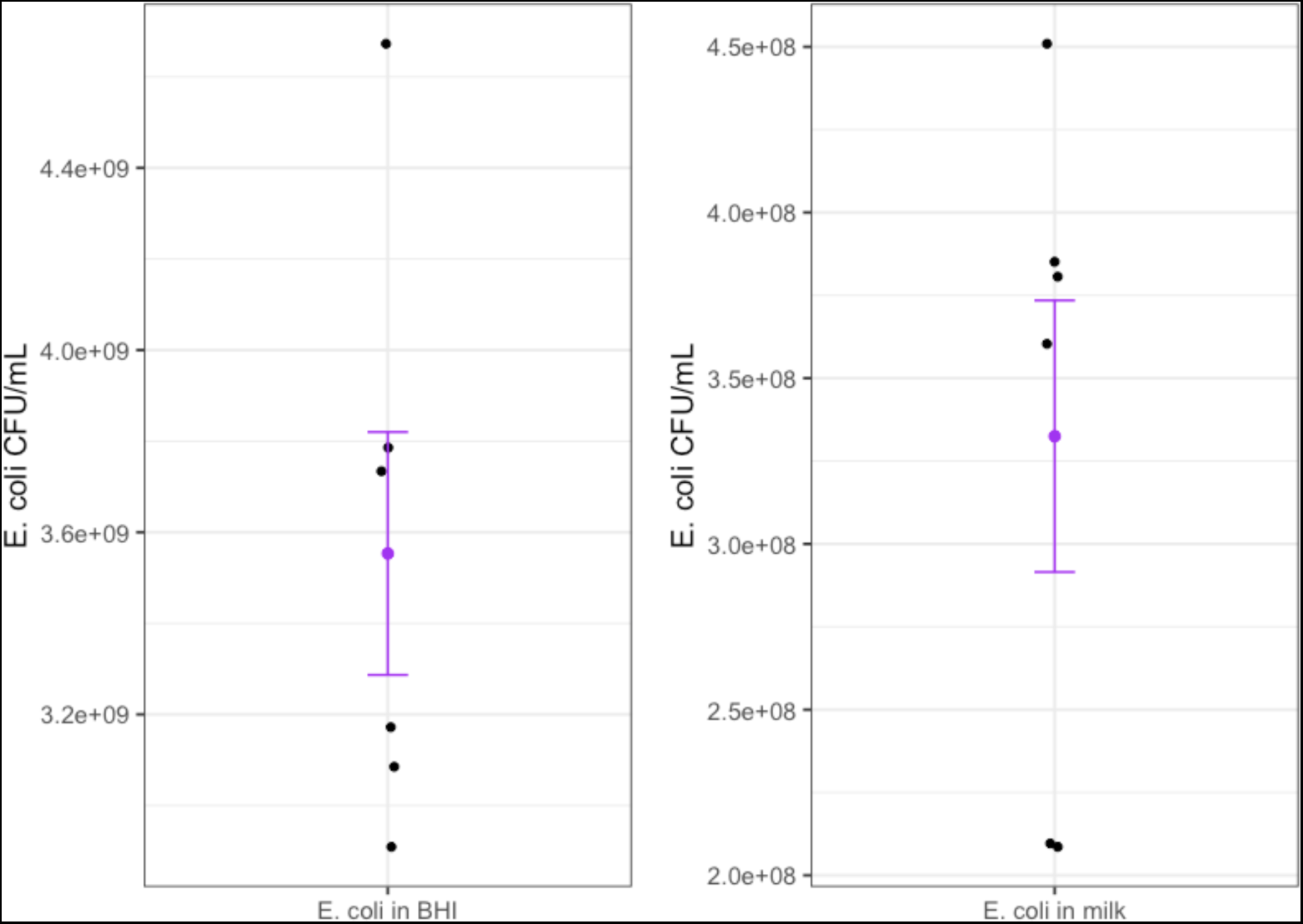
CFU/mL of *E. coli* growing alone for 72 hours in Brain Heart Infusion broth (37°C) and milk (28°C). Estimates calculated from colony counts on violet-red-bile (VRB) agar (replication = 6 biological replicates).

#### 5. DNA controls for Oxford Nanopore sequencing

<u>F1.1 control:</u> DNA isolated and previously sequenced in Leale et al. 2023

<u>M+ control:</u> mock synthetic community composed of DNA isolated from DSMZ strains and combined in attempted equal by DNA concentrations. *Lactobacillus delbreuckii (*DSM20072*), Lactobacillus helveticus (*DSM20075*), Acetobacter orientalis (*DSM15550*), Acetobacter lovaniensis (*DSM4491*), Limosilactobacillus fermentum (*DSM20052*), Lactococcus lactis (*DSM20481).

<u>Zymo control:</u> ZymoBIOMICS Microbial Community DNA Standard. *Listeria monocytogenes* – 12%, *Pseudomonas aeruginosa* – 12%, *Bacillus subtilis* – 12%, *Escherichia coli* – 12%, *Salmonella enterica* – 12%, *Lactobacillus fermentum* – 12%, *Enterococcus faecalis* – 12%, *Staphylococcus aureus* – 12%, *Saccharomyces cerevisiae* – 2%, and *Cryptococcus neoformans* – 2%.

**Figure S4:**
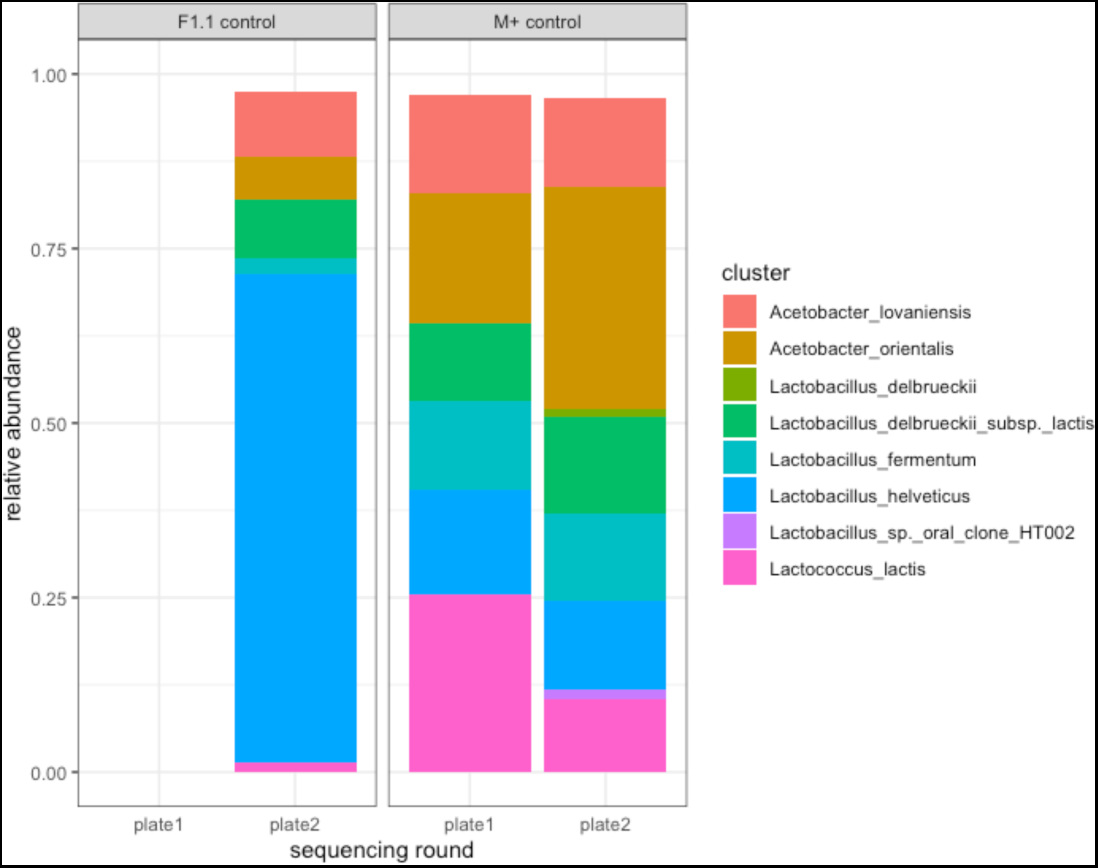
our sequencing and bioinformatic pipeline suitably distinguished bacterial types and their proportions predicted in Mabisi. Community compositions of Mabisi related positive control samples.

**Figure S5:**
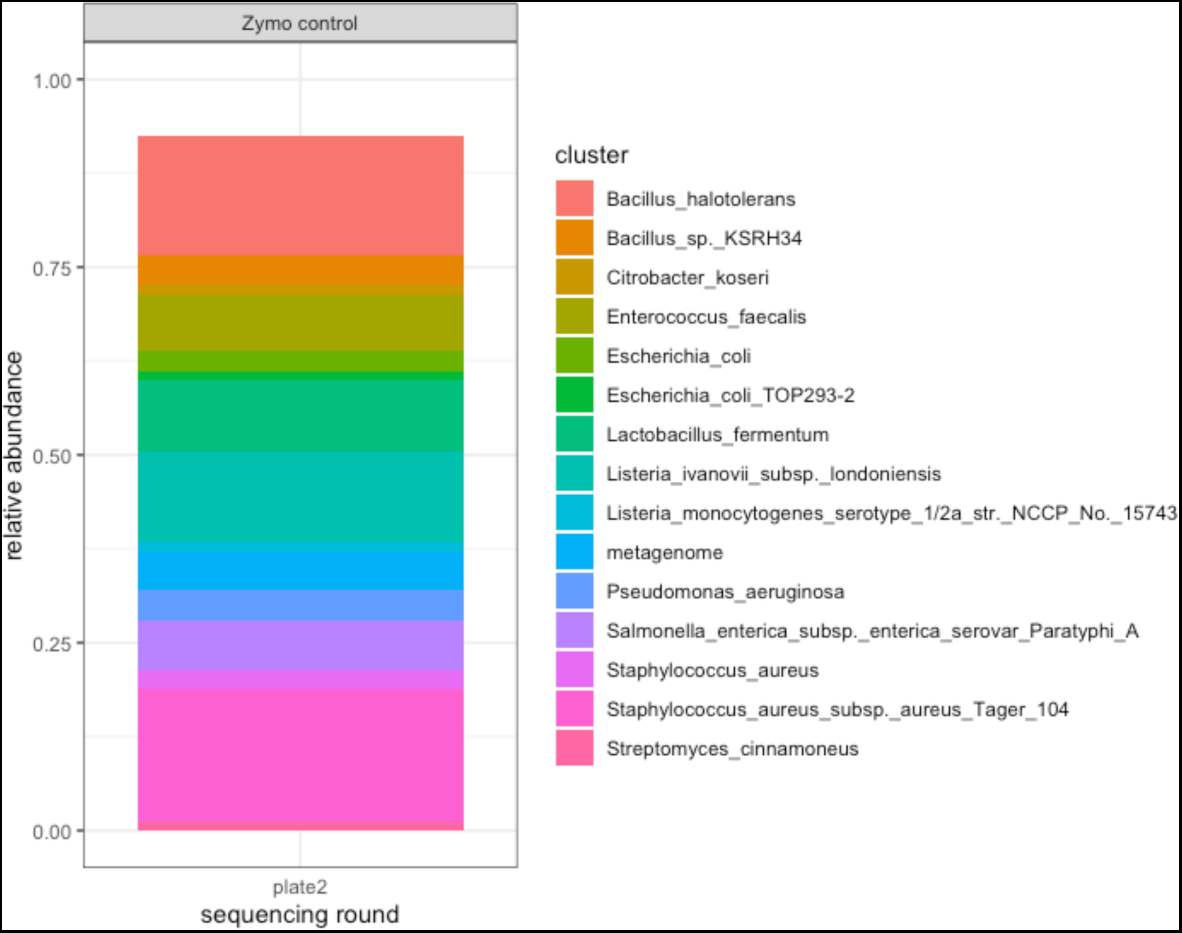
our sequencing and bioinformatic pipeline suitably distinguished bacterial types and their proportions predicted in Zymo postive control. Community compositions of ZymoBIOMICS Microbial Community DNA Standard.

